# Parallel genomic evolution of parasite tolerance in wild honey bee populations

**DOI:** 10.1101/498436

**Authors:** Katarzyna Bozek, Juliana Rangel, Jatin Arora, Mandy M.Y Tin, Emily Crotteau, Gerald M. Loper, Jennifer Fewell, Alexander S. Mikheyev

## Abstract

Sudden biotic pressures, such as those from novel diseases and pathogens, require populations to respond rapidly or face potential extinction. How this response process takes place remains poorly understood, particularly in natural environments. In this study we take advantage of unique decade-long data sets of two wild honey bee (*Apis mellifera*) populations in the United States to reconstruct the evolution of tolerance to a novel parasite, the ectoparasitic mite *Varroa destructor*. Upon the arrival of *Varroa*, the two geographically isolated populations simultaneously suffered massive *Varroa*-induced mortality, but stabilized within two years. Here we sequenced and phased genomes of 465 bees sampled from both populations annually over the decade that spanned *Varroa’s* arrival. Remarkably, we found that genetic changes were strongly correlated between the populations, suggesting parallel selective responses to *Varroa* parasitization. The arrival of *Varroa* was also correlated with an influx of genes of African origin into both populations, though surprisingly it did not substantially contribute to the overall similarity of the evolutionary response between them. Genes involved in metabolic, protein processing and developmental pathways were under particularly strong selection. It is possible that interactions among highly connected gene groups in these pathways may help channelize selective responses to novel parasites, causing completely unrelated populations to exhibit parallel evolutionary trajectories when faced with the same biotic pressure. Our analyses illustrate that ecologically relevant traits emerge from highly polygenic selection involving thousands of genes contributing to complex patterns of evolutionary change.

## Introduction

Identification of genes that underlie ecologically important traits, as well as the ability to link these genes to fitness effects, continues to be a central challenge in biology. Consequently, it remains unclear whether adaptation takes place via strong selection acting on a small number of genes with a large fitness effect, polygenic adaptation (Stinchcombe and Hoekstra 2008; Manolio et al. 2009), or genetic introgression from other populations (Olson-Manning et al. 2012). Most genome-wide association studies in both ecological and medical genomics rely on present-day data. Their statistical power is often limited, typically identifying relatively small numbers of loci with particularly strong fitness effects. Incidentally, such studies may also suffer from biases from population stratification and demographic history (Stinchcombe and Hoekstra 2008; Manolio et al. 2009). In contrast, historical studies that rely on long-term data collected throughout the course of a selective event can allow direct measurement of evolutionary change through time at every gene (Bi et al. 2013; Mikheyev et al. 2015; Yeates et al. 2016). Unfortunately, such historical data sets rarely exist, as the occurrence of selection events in the course of long-term field studies cannot be predicted, and proper collection requires considerable foresight and luck to adequately sample and preserve material for later analysis.

The emergence of novel pathogens that exert similar biotic pressure across multiple naive populations separated in space represents a unique opportunity to examine the course of evolutionary adaptation within a species. This opportunity has appeared in the Western honey bee (*Apis mellifera*), which has become host to an increasing range of parasites and pathogens since the late 20th century (Genersch and Aubert 2010). The ectoparasitic mite, *Varroa destructor*, represents a particularly severe example of a novel pest of honey bees introduced as recently as the late 1980s to the U.S. *Varroa* mites reproduce inside brood cells, feeding on larval and pupal tissues as well as adults, and acting as an efficient vector of several bee viruses (Ball and Allen 1988; Martin et al. 2012). *Varroa* mite infestations have caused sharp declines in honey bee populations around the globe (Finley et al. 1996; Martin et al. 1998; Vanengelsdorp et al. 2007). These declines have severely impacted management practices, as domesticated colonies typically require expensive and time-consuming annual treatments with acaricides for their continued survival (Francis et al. 2013). However, there are some feral honey bee populations that have been able to rapidly acquire tolerance to *Varroa*, often in just a couple of years after an initial period of high mortality (Pinto et al. 2004; Loper et al. 2006; Villa et al. 2008; Mikheyev et al. 2015). Studies of genome-wide historical changes in such populations can allow us to understand the mechanisms of resistance to *Varroa* as well as how selection for disease resistance occurs in natural populations (Behrens et al. 2011; Lattorff et al. 2015; Mikheyev et al. 2015).

Because of their agricultural and economic importance, honey bee populations are subject to intense monitoring and therefore well-sampled historical data sets exist (Mikheyev et al. 2015; Cridland et al. 2018). In particular, the U.S. Department of Agriculture had several monitoring sites of feral (wild) populations in the southern United States in the 1990s to track the projected arrival of Africanized honey bees (Pinto et al. 2005; Loper et al. 2006), which descended from African honey bees introduced to southern Brazil in 1957 (Kerr 1957). Africanized honey bees quickly spread northward over the following three decades, giving rise to aggressive hybrids infamously known as ‘killer bees’. Coincidentally, *Varroa* arrived during the same time frame. As a result, these time series studies provide an opportunity to examine how wild populations respond to novel parasites and pathogens. Furthermore, as Africanized honey bees are naturally resistant to *Varroa*, these data allow us to test relative contributions of major evolutionary forces, such as selection on standing genetic variation and immigration, to the evolution of parasite resistance.

## Results

### Demographic trends: parallel evolution of tolerance

In this study we compared the evolutionary responses of two unmanaged honey bee populations to *Varroa* parasitism. Each population was sampled yearly for a decade starting in the early 1990s, and the bees were stored frozen, minimizing post mortem DNA damage. The populations were located 1,400 km apart in starkly different environments, one in the Sonoran desert in Arizona, and the other in the coastal plains of Southern Texas (Fig. 1A) and were sampled during a period of relative climatic stability (Fig. S1). Initially the populations differed in subspecies composition, with the Arizona population consisting predominantly of colonies with Western European maternal ancestry (particularly *Apis mellifera mellifera*), while the Texas population consisted of colonies with mostly Eastern European maternal ancestry (*Apis mellifera ligustica*) (Loper 1995; Pinto et al. 2004). Coincident with the arrival of *Varroa* both populations experienced annual mortality rates tripling from 20% in 1990 to nearly 60% in 1995 (Loper 1995; Rubink et al. 1995). Remarkably, the populations recovered to approximately half of their original population size within two years and have remained at this strength until the present day (Fig. 1B) (Rangel et al. 2016). To characterize fine-scale genomic changes occurring as a result of novel parasite pressure, we sequenced and phased whole genomes from 465 individual worker bees (177 from Arizona, 288 from Texas), collected between 1991 to 2001, with between five to 40 individuals per year in each population. We found that the arrival of Africanized bees occurred just a few years after that of *Varroa* (Fig. 2), as reported previously (Loper et al. 1999; Pinto et al. 2005), raising the question of whether immigration could have been solely responsible for rapid evolution of mite tolerance in these populations.

**Figure 1.**
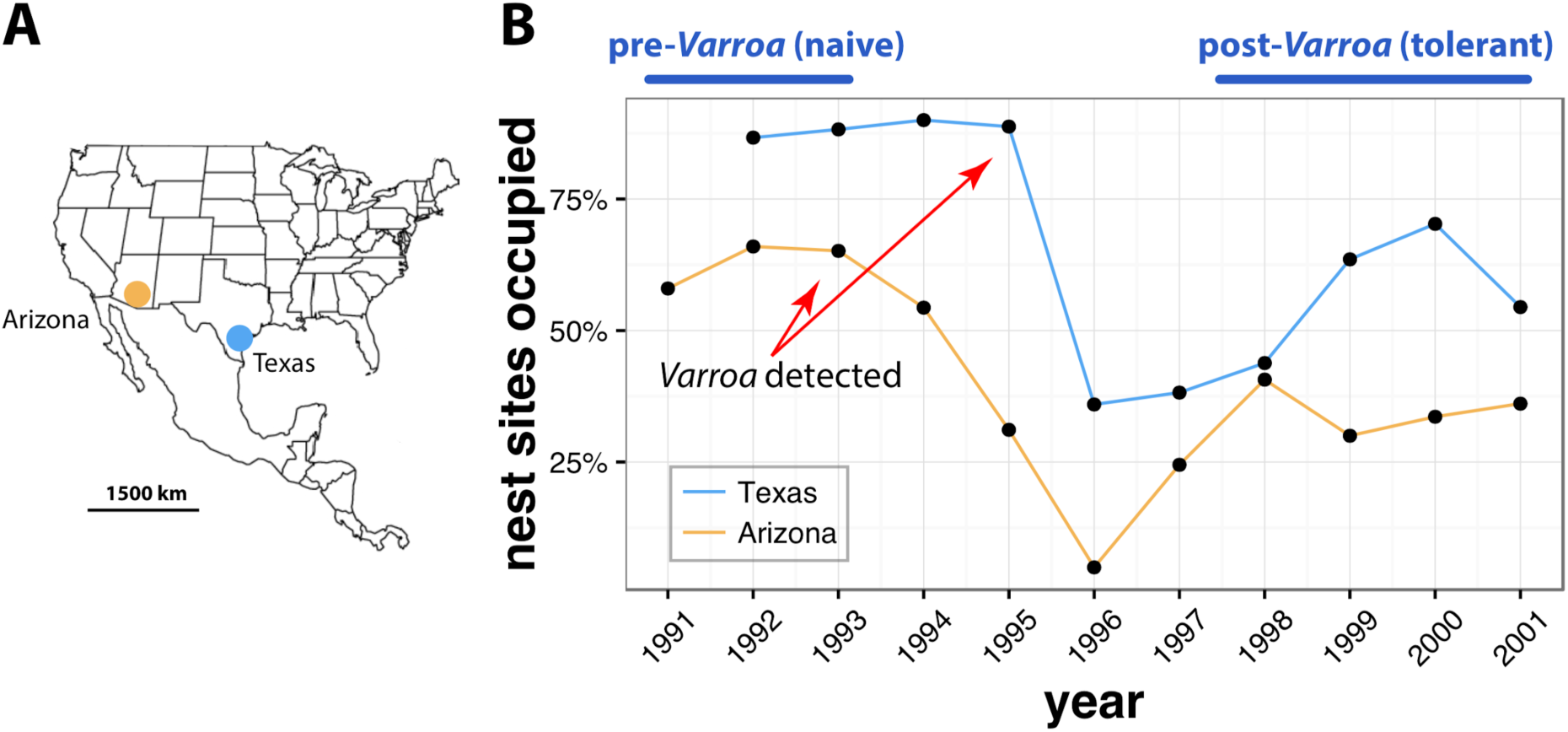
Population collapse and recovery after the arrival of *Varroa destructor* in two populations. (A) The two populations are located on the southern U.S. border. (B) Known nesting sites in two populations were sampled annually as part of feral and Africanized honey bee monitoring programs in Arizona and Texas. *Varroa* was first detected at a low level in 1993 in Arizona (1.4% of colonies infested increasing to 3.9% in 1994), and in 1995 in Texas, with both populations suffering heavy colony mortality in 1995–1996 and recovering to near *pre-Varroa* levels around 1998 (Loper 1995; Rubink et al. 1995). Subsequent surveys until 2010 in Arizona (this study) and a 2013 survey in Texas (Rangel et al. 2016), have shown that the populations have stabilized at the post-1998 levels. In this study we principally focused on comparing naive pre-*Varroa* populations (before 1994) with those that have evolved tolerance to this novel parasite (after 1997). Texas data below are from Table 1 in Pinto *et al*. (Pinto et al. 2004). Changes in population size were not liked to climatic factors, such as temperature and precipitation, which remained stable over the decade of observation (Fig. S1).

**Figure 2.**
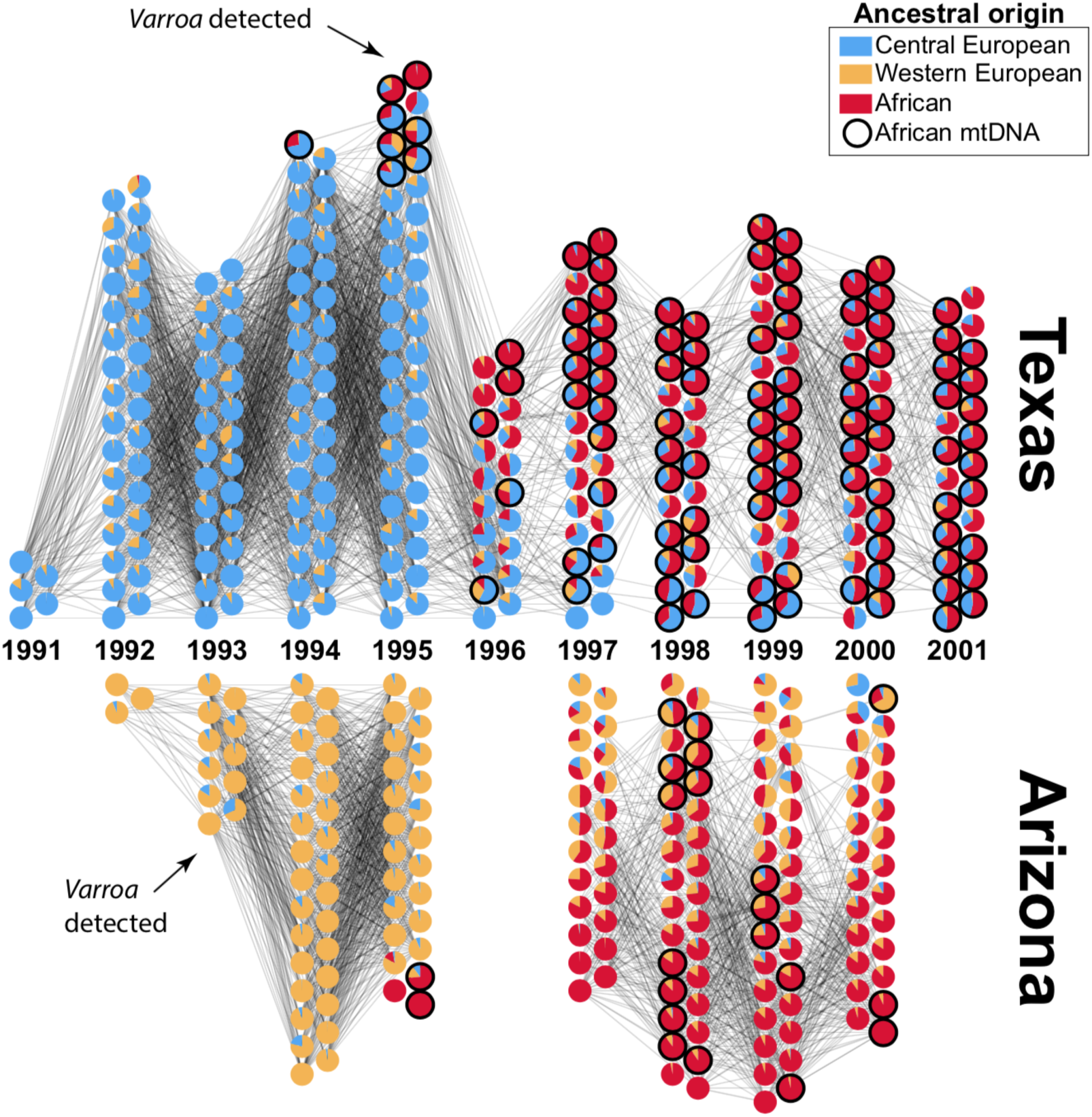
Immigration of Africanized honey bees preceded population collapses in 1996. Each circle represents an individual bee genome, partitioned into three possible ancestral populations (Central European, Western European, or African), with African mitochondrial haplotypes highlighted in red. Lines connecting individuals sampled in adjacent years correspond to 1% or more identity by descent. The arrival of Africanized honey bees preceded the *Varroa*-induced die-offs in 1996, and as evidenced by mitochondrial data, Africanization occurred by different mechanisms: through the establishment of new queens in Texas, and through male-mediated gene flow in Arizona. An interactive version of this figure is available at http://oist.github.io/bee-Varroa-visualization/

### Parallel genetic changes

We quantified microevolutionary changes in the populations over time by comparing changes in haplotype frequencies at each gene between naive and *Varroa*-adapted population using Wright’s fixation index (F_st_). Remarkably, changes in gene haplotype frequency after *Varroa* infestation were strongly correlated between the two populations (Fig. 3A). To test whether the observed F_st_ correlation was due to the selection specifically on genes, *vs*. on genome-wide architecture, we recalculated F_st_ for sequences with the same length distribution randomly sampled throughout the genome. If genes, rather than intergenic regions were the targets of selection, we expected to find higher F_st_ correlations associated with genic regions. This is indeed what we found (r = 0.54 ± 0.2 95% C.I. n simulations = 1000, *vs*. observed r = 0.83, Fig. 3A). The regions with correlated genetic changes were not close to each other in the genome, suggesting widespread evolutionary changes (Figures S2, S3). Likewise, haplotype frequencies of individual also changed in parallel in the two populations (Fig. 3B). This was true both of novel haplotypes, which were due to Africanized honey bee migration (Fig. 2), and pre-African haplotypes, though established haplotypes showed the greatest amount of phenotypic change (Fig. 3B). Interestingly, there was no overall loss of genetic diversity in most loci, despite the population bottleneck after the arrival of *Varroa* (Fig. S4).

**Figure 3.**
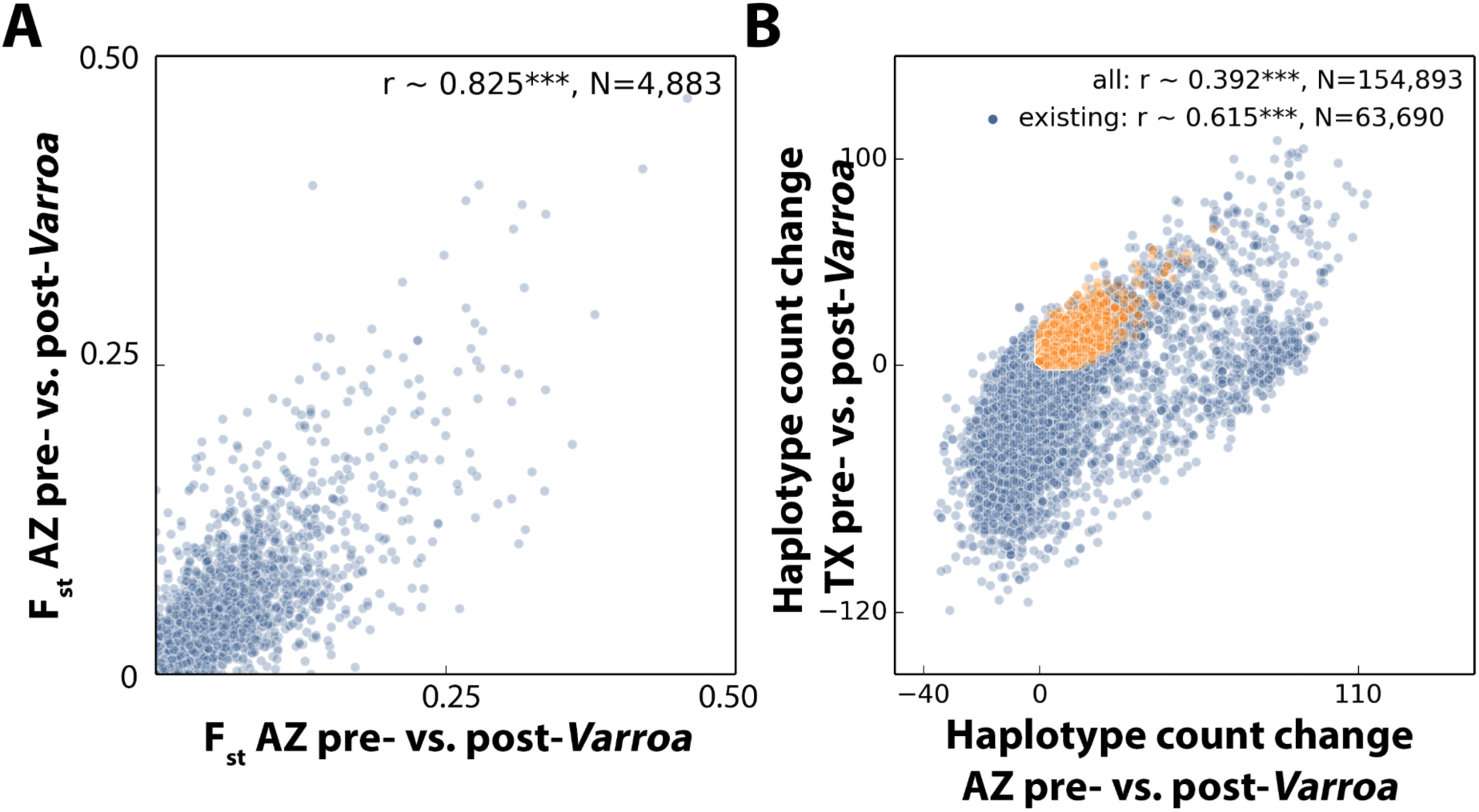
Arizona and Texas feral honey bee populations show parallel genetic evolution. (A) From before the arrival of *Varroa*, to the evolution of *Varroa* tolerance (Figure 1), 72% of the genes showed statistically significant changes in haplotype frequency in Arizona, and 82% in Texas based on F_st_ permutation tests. Genetic distances between pre- and post-selection populations were highly correlated, suggesting similar selective pressures operating at the scale of the entire genome. The F_st_ score was, however, independent of gene location on any chromosome (Fig. S2). Because absolute measures of distance are sometimes better than F_st_ for differentiating evolutionary scenarios (Cruickshank and Hahn 2014), we re-calculated this analysis using correlations of d_xy_/d_x_ and obtained essentially the same result (N = 4883, r = 0.84). (B) In addition to the parallel changes at the level of summary statistics (F_st_), counts of haplotype frequencies for each gene also changed in the same direction (Figure S10 shows a breakdown by chromosome).

### Functional characterization of genetic changes

Given the large number of genes involved, comprehensive functional characterization is difficult. However, genes that showed the greatest shift in F_st_ in pre- and post-Varroa within populations were enriched for similar functional categories and pathways between populations (Tables S1 and S2). Genes associated with metabolism, gene expression, protein regulation, and signal transduction gene ontology (GO) terms showed particularly strong changes over time (Table S1). Many of these genes belonged to specific biochemical pathways, which allowed for further functional characterization of the changes over time (Table S2). This included genes involved in signal transduction belonging to Hippo and Notch signalling pathways, which are important in development, as well as metabolic and ion transport genes in the citrate cycle and oxidative phosphorylation pathways. Given the pleiotropic nature of these pathways, identifying the precise mechanisms of tolerance is challenging. However, large-scale changes in development could have an effect on the reproductive rate of *Varroa* on worker brood, particularly given the reduction in worker size post-*Varroa* (Fig. S5) (Mikheyev et al. 2015).

### Immigration *vs*. selection for *Varroa* tolerance as the driver of parallel changes

Selective pressures on natural populations may act on both standing genetic variation and beneficial alleles from immigrants (Hedrick 2013). Both of these mechanisms could have been at work in Arizona and Texas, given the massive gene flow from Africanized bees during the time of this study (Fig. 2) (Loper et al. 1999; Pinto et al. 2004; Pinto et al. 2005; Harrison et al. 2006). Thus, either similar selective pressures or immigration could account for the observed parallel genomic changes in both populations, as well as for the parallel phenotypic shifts, such as the reduction in worker size characteristic of Africanized honey bees (Camazine 1986; Nunes-Silva et al. 2006; Francoy et al. 2008) (Figures 1 and 2, Fig. S5); these effects need not be mutually exclusive, however. We performed a number of tests in an attempt to disentangle the role of immigration from selection, in particular focusing on the null hypothesis that the observed pattern can be solely attributed to Africanized immigration, which appeared to be the dominant signal in the data at first glance (Fig. 2).

First, we conducted a partial regression on the correlation between pre- and post-*Varroa* F_st_ changes within population, using the F_st_ between post-*Varroa* population as a covariate. The rationale for this analysis assumes that, if most similarities in the post-*Varroa* populations are due to shared African heritage, the covariate should explain much of the variation in the between-population correlation across times. However, it explained only a relatively minor percentage of the total variation in the correlation of F_st_ changes between populations, suggesting that most correlated changes were not due to shared ancestry (partial R^2^ = 0.90%).

Second, we also calculated F_st_ in the pre- and post-*Varroa* infected populations after filtering out haplotypes that emerged after 1994, keeping only those present before any African genetic admixture was detectable, and before *Varroa*-induced mortality took place (Fig. 2). After haplotype filtering, F_st_ distance between post-selection populations was higher than between the pre-selection populations, suggesting that genes of African descent were successfully removed using this approach (Fig. S6). Despite a reduction of statistical power caused by the elimination of rare haplotypes, the filtered F_st_ distances between pre- and post-*Varroa* populations remained significantly correlated (Fig. 3B). Furthermore, genes that received more immigrant (putatively Africanized) alleles, showed significantly lower F_st_ correlations between populations, suggesting parallel genetic changes were not driven by the immigrant alleles (r = ×0.73, n = 4,890, p < 2 ×10^−16^).

Third, we quantified the relationship of genetic changes in the populations with their mortality and phenotypic data using Granger causality. This metric examines whether one sequence of time series data carries information to predict another. We found that in all cases, the extent of Africanization was not predictive of changes in nest site occupancy, mortality and recovery (Wald test, 1 d.f., p > 0.5 for both populations). However, mortality was predictive of the arrival of Africanized bees in Arizona (Wald test, 1 d.f., p = 0.0013), though not in Texas (Wald test, 1 d.f., p = 0.76). Thus, increased nest site availability in Arizona might have facilitated immigration of Africanized bees following the *Varroa*-induced population collapse.

Because a reduction in body size was previously hypothesized as a possible adaptive response to *Varroa* tolerance (Mikheyev et al. 2015), we examined whether there was such a change in the studied populations. As Africanized bees are smaller than European bees, we further investigated whether any size effect may have been driven by Africanization. Individuals in both populations were indeed smaller after the arrival of *Varroa*, although the trend for smaller size in Arizona started well before the arrival of Africanized bees, while in Texas there was a more abrupt transition of both size and Africanization patterns (Fig. S5). Thus, Africanization was not necessary for body size changes, although it could have been a contributing factor.

Finally, we used approximate Bayesian calculation to examine whether observed correlation in haplotypic F_st_ values could be due purely to between-population migration or immigration from the same large Africanized population. We created a forward-time simulation that closely mimicked the biology and genetic structure of the bee populations under study. Within this framework, we found decisive support of the model with parallel selection within populations, as opposed to a migration-only model (Bayes factor = 494). The resulting summary statistics closely resembled the observed data (Fig. S7).

## Discussion

The observed evolutionary responses in the two honey bee populations were highly polygenic (Fig. 3), and were most likely a result of convergent selection regimes. Surprisingly, given the striking influx of Africanized bees in Arizona and Texas during this period (Fig. 2), immigration seems to have played only a partial role in the evolution of *Varroa* tolerance, based on the extensive analyses we conducted. Notably, this finding matches the rapid evolution of resistance in another USDA-monitored population in Louisiana, where *Varroa* was first detected in the same time frame (early 1990s), but was outside the geographic range of Africanized bees until the mid-2000s (Villa et al. 2008). Strikingly, the Louisiana population rebounded to pre-*Varroa* levels within five years. Unfortunately, no biological samples were preserved, so we were unable to run similar analyses on this population. One possibility for the surprisingly minor role of Africanization on the rapid adaptation of the bee populations to *Varroa* parasitism is that, for introgression to facilitate adaptation, beneficial traits need to be expressed in the F_1_ hybrid (Hedrick 2013). But even though Africanized bees are generally more resistant to *Varroa*, any of their brood that hybridizes with European bees may actually be more susceptible to the mite (Guzman-Novoa et al. 1996), limiting short-term benefits of hybridization. Thus, pre-existing genetic variation might play a more important role than immigration in the evolution of *Varroa* tolerance.

With the caveat that correlational studies, particularly in the field, cannot account for all possible factors at play, a major strength of the current study is that it captures year-to-year changes in unmanaged populations in response to a novel parasite and associated zoonotic diseases. Consistent with current knowledge of *Varroa* impact on honey bees, both populations showed marked reductions in population size after the mite arrival (Fig. 1), which acts as potent selective agent of honey bee populations (Lattorff et al. 2015; Mikheyev et al. 2015). While *Varroa* tolerance was not directly assayed over time, the demographic bottleneck and recovery shown in Figure 1 indicates that the bees were at first sensitive to *Varroa* but soon evolved resistance at the population level. This has been reported previously (Villa et al. 2008), and is indicative of strong selection and its response. We were also able to rule out neutral causes such as migration, immigration and drift as sole drivers of these large-scale genetic changes (Fig. S7). Thus, it seems likely that parallel evolution was driven by *Varroa*, though strictly speaking, we cannot rule out other causes during the period of the same few years.

That being said, our data demonstrate that ecologically relevant evolutionary changes, such as the evolution of tolerance, can be associated with wide-ranging, genome-wide changes that are consistent across populations. These findings are consistent with the growing body of literature that has shown parallel genetic and genomic changes in both natural and experimental populations of sexually reproducing animals (Colosimo et al. 2005; Graves et al. 2017; Nosil et al. 2018). Phenotypic and genomic parallelism can be particularly prevalent under ‘soft sweep’ conditions when population sizes are large (Karasov et al. 2010), which is especially true of insects. Similar selective regimes may explain the remarkable resistance of many natural populations to biotic and abiotic stressors, and the widespread occurrence of parallelism in nature.

### Materials and Methods

#### Surveys of feral honey bee populations

Analysis of genetic changes over time is limited by the availability of well-sampled, comparable and high-quality historical material. Fortunately, samples from two long-term study sites set up by the U.S. Department of Agriculture (USDA) in Arizona and in Texas survived to the present day as part of its program to monitor the arrival of Africanized honey bees, which were expected to arrive in the 1990s. The arrival of *Varroa* occurred unexpectedly during this time, and was noted by the USDA researchers. In Arizona, from 1989 to 1991, feral honey bee colonies were discovered along Camp Grant Wash and Putnam Wash in Pinal Co. in caliche outcroppings following a series of washes along an approximately 1 × 15 km^2^ area. A total of 245 colonies were discovered and mapped using GPS coordinates, and each colony location was given an identification name and number. Accessible colony sites were resampled seasonally (generally March, June and October) each year, and not all colonies were accessible for genetic sampling. Surveys were conducted annually from 1991 to 2001, then sporadically in 2003, 2004, and 2010 (Loper et al. 1999; Harrison et al. 2006; Loper et al. 2006). In Texas, a population of feral honey bees was sampled annually from 1991 to 2001, and again in 2013. Texas survey methods have been described previously (Pinto et al. 2004; Rangel et al. 2016). Specimens were collected on dry ice in the field, and stored at −80°C.

#### DNA library preparation and sequencing

The forelegs of one honey bee per colony per sample year were used for DNA extraction. They were placed into a 96-well U-bottomed titer plate (Greiner Bio-One) containing 4 mm stainless steel beads in each well (Taitech) and homogenized in 200 μl extraction buffer as described in Mikheyev *et al*. (4) with 200 μg/ml proteinase K (Nacalai Tesque, Inc.) at 1000 rpm for 30s. The tissues were homogenized again for 30s after a 30s rest and incubated at 56°C overnight. DNA was extracted using silica-magnetic beads following Mikheyev *et al*. (4), but here the whole process was automated on a BioMek FXp platform (Beckman Coulter) (Tin et al. 2014). DNA concentration was measured by Quant-iT PicoGreen dsDNA assay kit (Invitrogen). Individually barcoded Illumina Nextera XT libraries were then prepared using the manufacturer’s protocol, with normalization and size selection performed on a BioMek FXp platform, as described in Mikheyev et al (2015). Libraries were sequenced on an Illumina HiSeq 2000 instrument by BGI in paired-end mode for 100 cycles from each fragment end. In total, we sequenced 465 diploid worker genomes at a depth of 7.2 ± 2.3 (S.D.) and 19 haploid drone genomes at a depth of 20.6 ± 3.0.

#### Alignment and variant calling

Using bowtie2 (2.2.6) (Langmead and Salzberg 2012) in “very sensitive” mode, we aligned reads to the Amel_4.5 genome reference (Honeybee Genome Sequencing Consortium 2006), and separately to the mitochondrial genome (Crozier and Crozier 1993). Genotypes were called using Genome Analysis Toolkit (GATK, 3.3), freebayes (1.0.2), and samtools (1.2) (Li et al. 2009; McKenna et al. 2010; Garrison and Marth 2012). Variant calls were converted to allelic primitives using GATK and combined using BAYSIC (Cantarel et al. 2014), keeping only those sites with at least 0.80 posterior probability. Additionally, sites where drones were found to be diploid were removed, as they were considered likely artifacts. Aligned and filtered results were sorted and separated by chromosome number.

#### Orienting scaffolds

The Amel_4.5 reference genome has several scaffolds that, while assigned to linkage groups, are not oriented with respect to the other scaffolds. In an effort to improve imputation accuracy, we used patterns of linkage disequilibrium decay, as measured by r^2^ and calculated using VCFTools (Danecek et al. 2011) to orient the scaffolds in the same order. We then modified the annotation to correspond to the newly oriented coordinate system. The new genome and annotation are available in the accompanying code repository on GitHub.

#### Genome phasing and validation

Reads from nine pairs of drones were sub-sampled to the mean worker coverage and mixed together to create ‘synthetic diploid’ individuals, which were added to the worker samples prior to imputation. These samples were then used to test the quality of imputation through comparison to the original haploid drone samples, for which phase was known, and which were sequenced at a higher depth. We next tested several cutoffs allowing for at most 50, 40, 30, 20, 15, 10, 5, or 1% of missing sequence data in diploid samples for any given position. Phasing and imputation were performed on the data limited this way using BEAGLE (4.1) (Browning and Browning 2009). Switch error and amount of missing imputation were assessed for different missing-data cutoffs by comparing the synthetic diploid genomes with the pre-imputation initial haploid drone sequence data. Switch error was computed using VCFTools, while missing data imputation was calculated based on the comparison of the matching phased and unphased vcf file content. Accordingly, we chose 10% of missing sequence data as a cutoff, which resulted in minimal switch error (0.0093 on average) and imputation loss (0.0641 on average), retaining 1,443,281 of the initial 10,147,753 SNPs (Figures S7 and S8).

#### Ancestral population assignment

Prior work has shown that the two study populations were largely descended from Eastern European (*A. m. carnica* and *A. m. ligustica*), Western European (*A. m. mellifera*), or African bees (*A. m. scutella*) (Loper et al. 1999; Pinto et al. 2004; Schiff et al. 2004; Rangel et al. 2016). As a result, we ran ngsAdmix (Skotte et al. 2013) with three ancestral populations using nuclear genetic data. The mitochondrial genetic distance matrix was used to classify mitochondria as being either of African or European descent with a k-means classifier (k = 3) using previously published data as a training set (Harpur et al. 2014; Wallberg et al. 2014).

#### Genome-wide changes within and between populations

Genetic change was quantified as F_st_ scores calculated for each population by comparing pre- and post-*Varroa* time periods using PopGenome (Pfeifer et al. 2014), using both coding and non-coding sequence in the most current genome annotation (Elsik et al. 2014). For each gene we computed the significance of the F_st_ score using permutation. Since most genes showed statistically significant frequency shifts, we also computed correlations in the F_st_ changes between the Arizona and Texas populations. To determine whether the observed changes were unique to genes, we repeated this analysis on non-gene sequences randomly sampled across the phased genome. For each gene we randomly selected a location in the genome and from every individual’s genome we extracted sequences of the same length as the respective gene. This way the randomized non-gene data had the same volume as the respective gene data. F_st_ scores were estimated in the same way as in the gene sequences. Confidence intervals for each correlation were bootstrapped using 1000 pseudo-replicates.

Although the primary analysis was based on haplotypic scores, we confirmed that qualitatively similar results were correlated with nucleotide F_st_ using datasets before and after haplotype imputation procedure. We also cross-checked our results by repeating the correlation analysis using an absolute measure of sequence divergence (d_xy_ divided by pre-*Varroa* d_x_).

#### Estimating the role of immigration

We employed three independent approaches to distinguish whether genetic change in the Arizona and Texas populations was due to selection *vs*. re-colonization by Africanized immigrants. The first approach employed a linear model comparing distances between populations of the form F_st_(AZ_pre_AZ_post_) ~ F_st_(TX_pre_TX_post_) + F_st_(AZ_post_TX_post_), where F_st_(XY) represents F_st_ score between populations X and Y, AZ_pre_, TX_pre_ represent Arizona and Texas populations before 1995, respectively, and AZ_post_, TX_pos_ represent Arizona and Texas populations from 1998 onward. The cutoffs were chosen based on prior studies, which found that allele frequencies had stabilized in those populations by 1998 (Pinto et al. 2005; Harrison et al. 2006). Specifically, the first term measures the extent of the parallel selective response across populations, while the last term models the degree of shared ancestry in the post-selection populations, presumably due to immigration from a common ancestral population. We then compared the model fit with and without the F_st_(AZ_post_TX_post_) term.

In the second approach, we excluded all haplotypes that were not detected prior to *Varroa-*induced population collapse (from 1995 onward), and large-scale African genetic admixture (Loper et al. 1999; Pinto et al. 2004). We then re-computed haplotypic F_st_ scores between populations in the filtered data sets.

In the third approach, we used Granger causality to determine whether mortality, as measured by site occupancy, contained any information that predicted the amount of African genetic admixture over time. This test specifically measures the ability to predict the future values of a time series with past values of another time series. We chose a lag of one year for the analysis, assuming a tight coupling between mortality and Africanization.

#### Africanization rates of individual genes

The test of filtering out new potentially Africanized haplotypes described above was conducted to measure the extent of per-gene Africanization. In such test, the amount of Africanized gene flow for every gene was estimated as the fraction of haplotypes not found in the population prior to 1995, before Africanization became prevalent (Fig. 3). This metric allowed us to test whether genes with a greater percentage of African contribution were more or less likely to show correlated genetic shifts over time, *e.g*, because African alleles are incompatible with European alleles.

#### Approximate Bayesian Computation (ABC) comparison of evolutionary scenarios

We used simuPOP (Peng and Kimmel 2005) forward-time population genetics simulation environment together with Approximate Bayesian Computation (ABC) (Csilléry et al. 2012) to estimate whether within-population selection was a significant contributor to F_st_ correlations between populations, as opposed to Africanized immigration or migration between the two populations. The simuPOP package is particularly suitable as it allows models including haplodiploidy and polyandry. The source code for the simulation is available in the online code repository and we summarize it briefly below. We simulated the evolution of Arizona and Texas populations over ten generations, with empirically observed population size changes (Fig. 1). We assumed that the Africanized population, which spanned much of Mexico and the southern U.S., was much greater than the target populations and remained at a constant size. We assumed that Africanized immigrants were highly resistant to *Varroa*, and that resistance was an additive polygenic trait. The simulation had four estimated parameters: (1) positive selection on the resistance alleles (initiated at the observed minor allele frequency in the Arizona and Texas populations), (2) immigration from the Africanized population, (3) migration between Arizona and Texas populations, (4) initial size of the Arizona and Texas populations. Onset of effects of (1) and (2) was set at generation five, equivalent to the estimated time arrival of *Varroa*. This framework allowed us to specifically compare scenarios with and without selection in order to estimate whether the observed patterns could be due to migration alone. For each scenario we generated 25,000 simulations each one containing 100 genes randomly sampled at empirical frequencies from the data set, computing F_st_ correlations from numbers of individuals equal to the number in our dataset randomly sampled from the first and last population in the simulation.

#### Changes in wing size over time

Nineteen landmarks at vein junctions (Francoy et al. 2008) were photographed under a Nikon SMZ1500 microscope, and digitised using the CLIC package (Dujardin) with a Wacom Intuos tablet. Procrustes analysis was performed on raw coordinates and centroid sizes of all samples were extracted using Shapes package (Dryden 2015). Centroid sizes were averaged for multiple wing measures. We then estimated the effects of African admixture, collection year and population on body size using linear regression.

#### Functional analysis

We used existing annotation from SwissProt as well as results of Annocript (Musacchia et al. 2015) for predicting gene function. Out of 15,289 genes reported in the bee genome we could associate GO terms to 8,177 of them and KEGG pathways to 1,345. For every GO term or KEGG pathway assigned to at least ten genes, we evaluated its relative contribution to F_st_ using a linear model including each functional term as a predictor, and the F_st_ distance between post-selection populations (as a covariate controlling for African genetic admixture), with no intercept. This analysis was carried out separately for Arizona and Texas populations. The confidence interval of each coefficient was determined using nonparametric bootstrap with 10,000 pseudoreplicates.

#### Source code availability

Source code used in genomic component of this project can be found at https://github.com/oist/varroa_parallel. The R script and results main statistical analyses can be found at https://oist.github.io/varroa_parallel/stats.html.

#### Data accessibility

Raw genomic reads were deposited to DDBJ and are available as BioProject PRJDB3873.

## Acknowledgements

We are grateful to Martin Helmkampf for assistance in shipping bees, and to Miquel Grau-Lopez for help with the analysis. Funding for this work has been provided by the Okinawa Institute of Science and Technology Graduate University.

## Author contributions

Designed the study: ASM; Performed field work: GML; Contributed specimens: JR, JF; Performed laboratory work: MYYT; Analyzed data: KB, ASM, JA, EC; Wrote the manuscript: ASM and KB.

## Supplemental Material

### Supplemental Figures

**Fig. S1.**
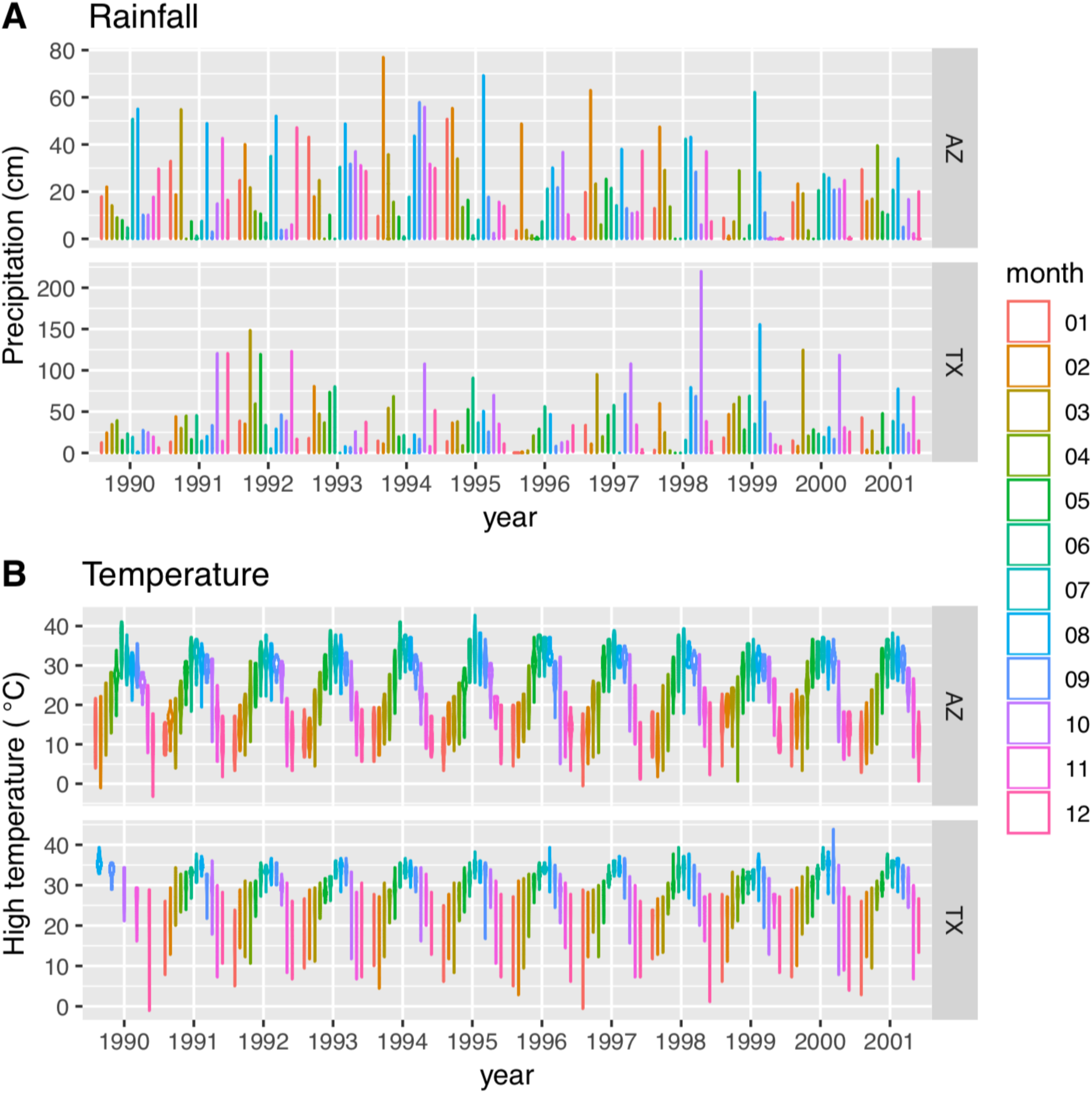
Climatic patterns did not notably change over the course of the observation period. Climate data were collected from the actual Texas field site, and from closest weather station in Arizona (Oracle, AZ) (retrieved from https://www.climate.gov).

**Fig. S2.**
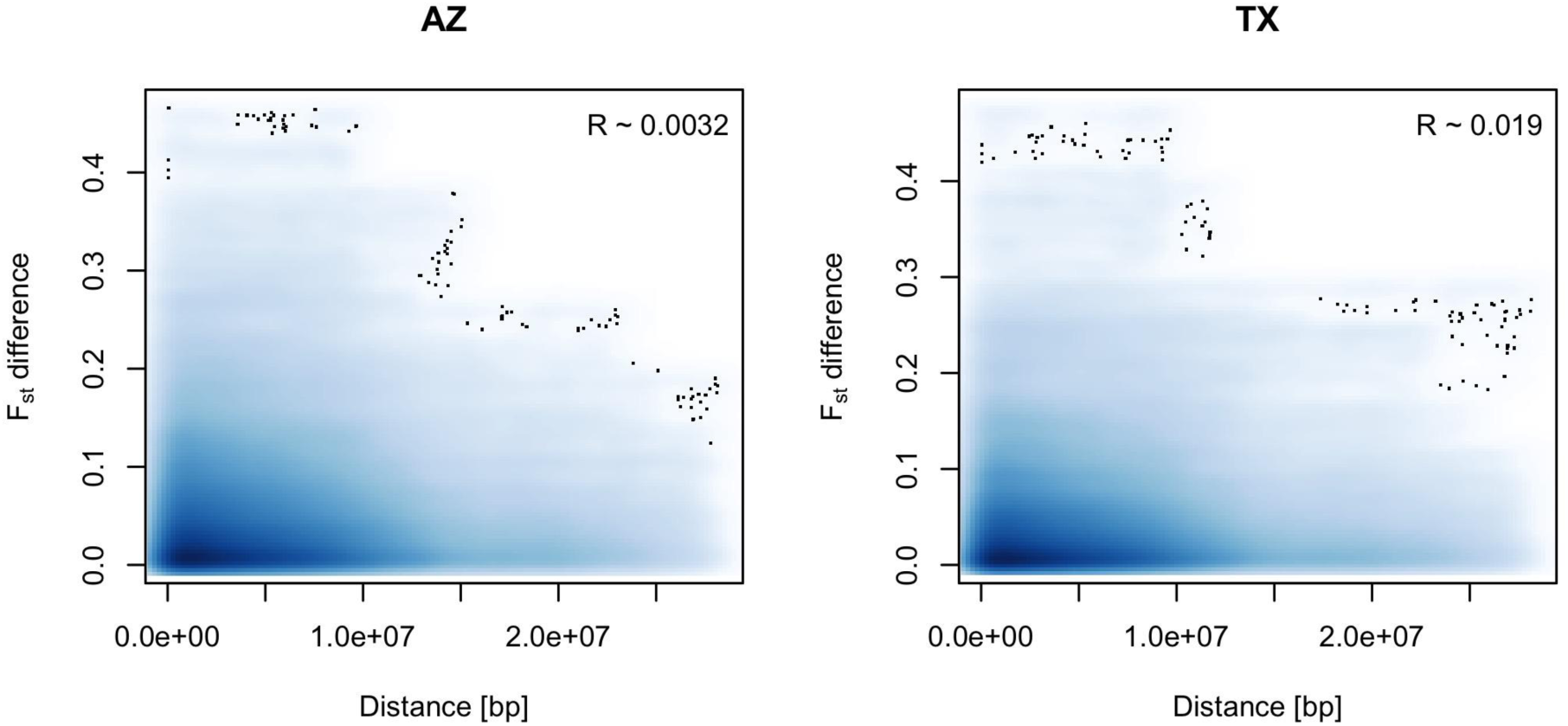
F_st_ differences vs. chromosome location. Neighboring location on a chromosome does not indicate similar F_st_ scores. Distance among genes in base pairs was compared to the absolute difference in the F_st_ score for all pairs of genes located in the same chromosome. Higher concentration of points is indicated by darker shades of blue and outliers are marked with black dots.

**Fig. S3.**
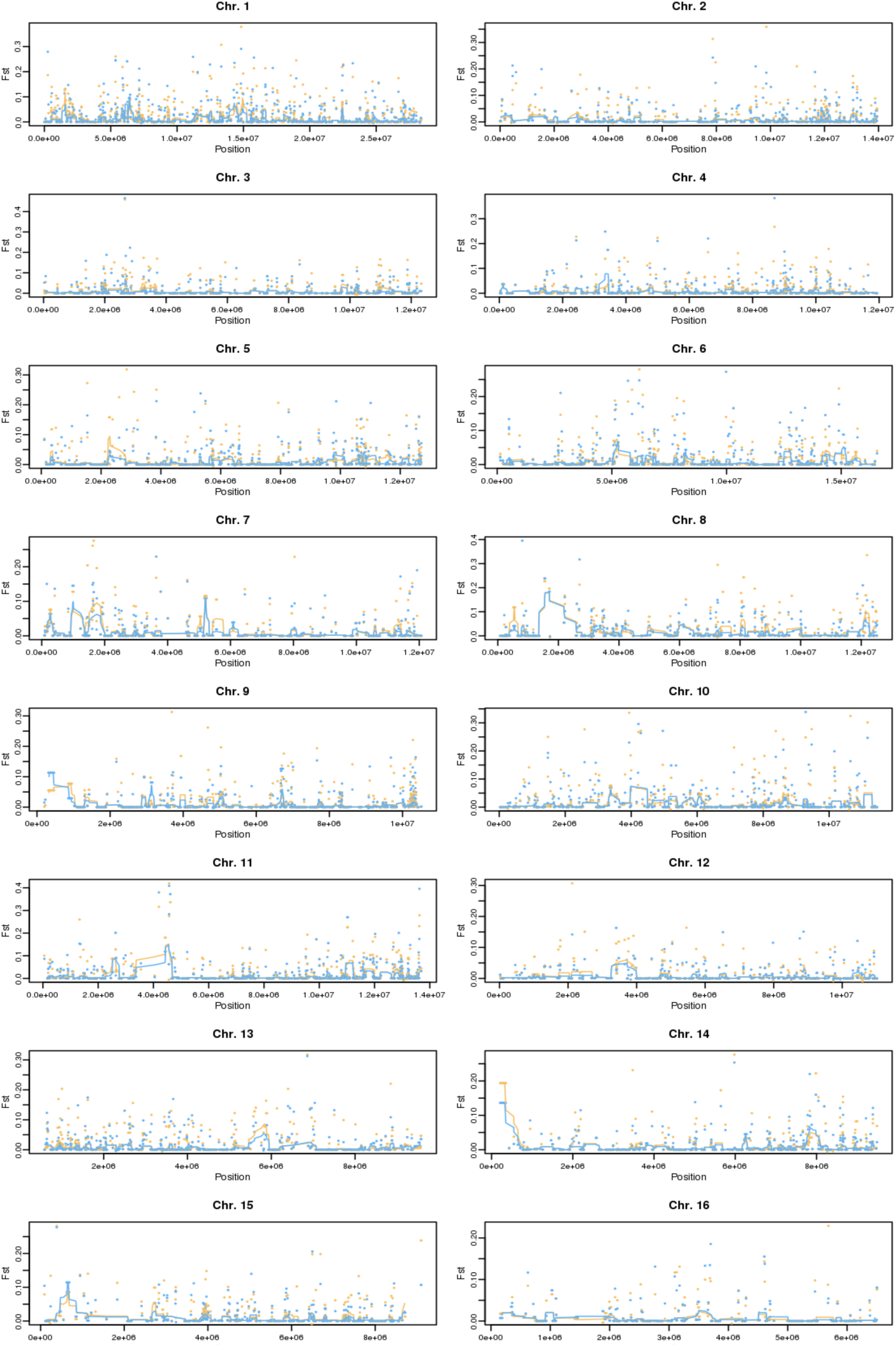
Manhattan plot of F_st_ values for individual genes and lines corresponding to a 50 kilobase rolling average. See interactive version at https://oist.github.io/varroa_parallel/manhattan.html.

**Fig. S4.**
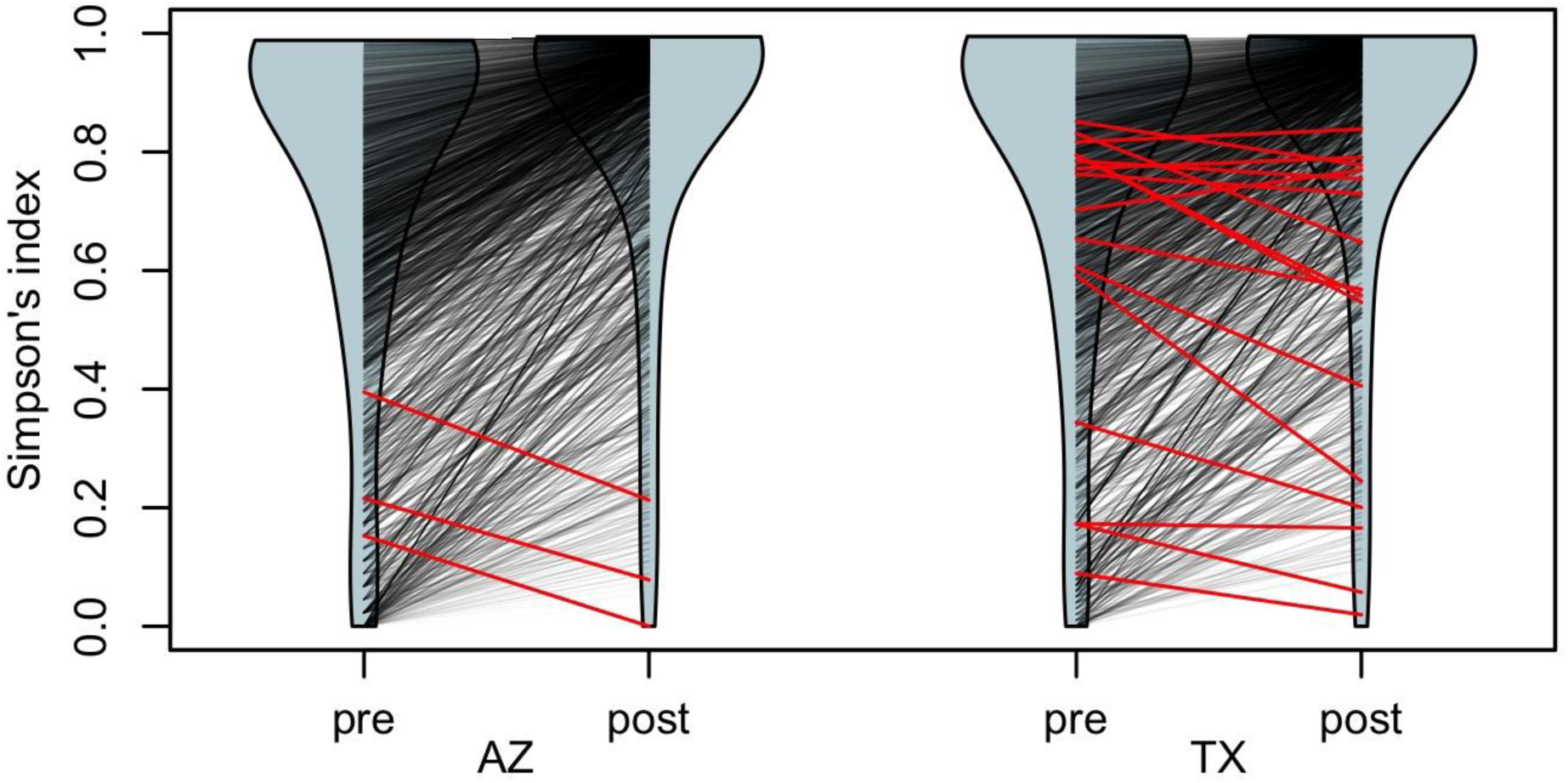
Haplotype diversity index. Haplotype diversity assessed using Simpson’s index in both populations shifted predominantly to higher values in the populations after *Varroa* exposure. The violin shapes represent diversity index distribution before (pre) and after (post) exposure to *Varroa* in the Arizona (AZ) and Texas (TX) populations. Black lines connect matching genes in the pre- and post-*Varroa* distributions, while red lines indicate genes that underwent a significant decrease in diversity after the arrival of *Varroa* (3 genes in AZ and 19 genes in TX). Many genes increased in diversity likely as a result of Africanized honey bee immigration into these populations.

**Fig. S5.**
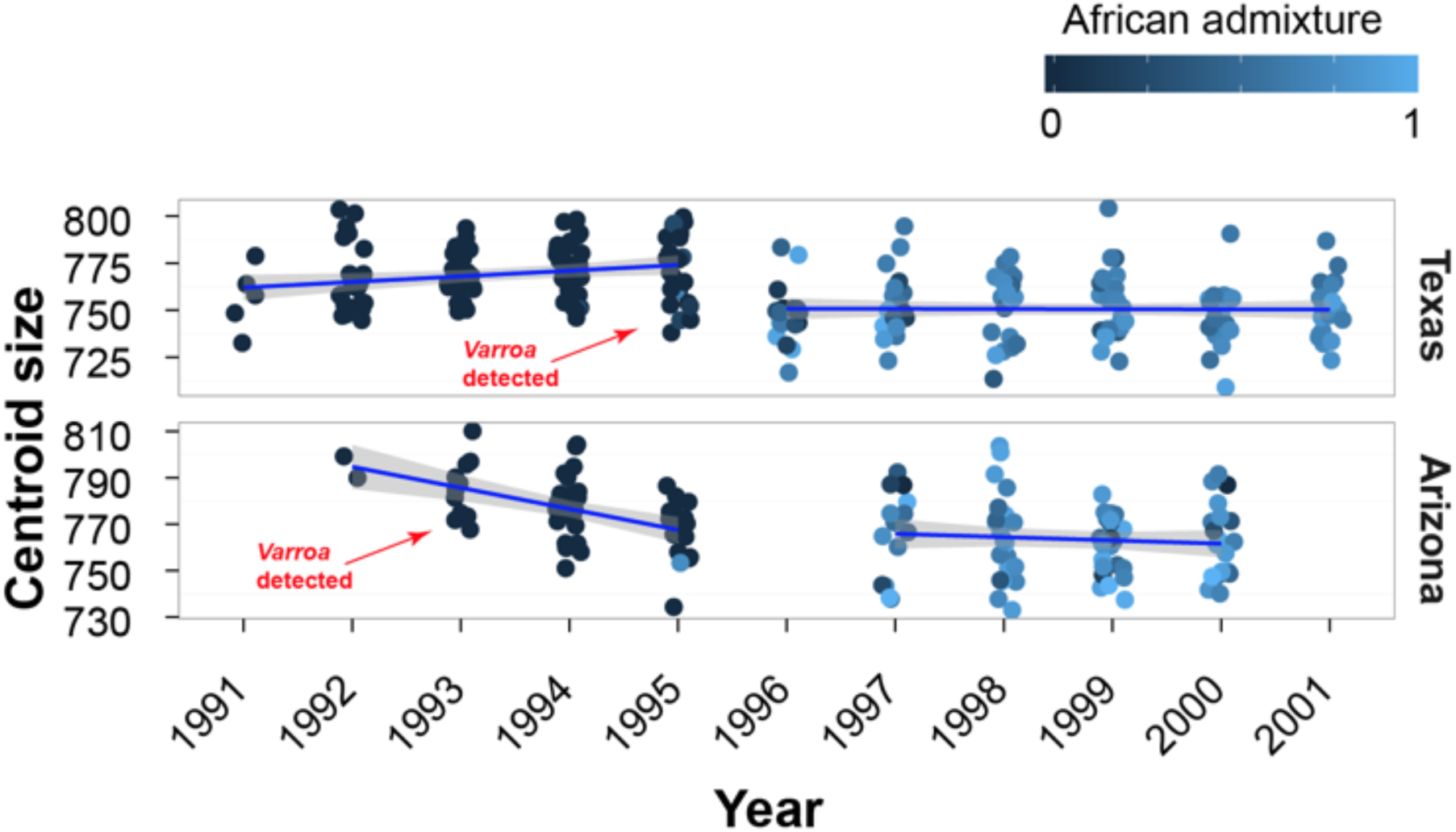
Size of honey bee workers reduced after the arrival of *Varroa*. Points jittered around years correspond to samples from individual colonies, with lighter colors corresponding to the increased African genetic admixture (see also Fig. 3). The arrival of *Varroa* and Africanized bees in 1996 corresponded to a decrease in average worker size. After 1996, the extent of African genetic admixture explained a small but significant proportion of variation in worker wing size (linear regression partial R^2^ = 2.9%, p = 0.02).

**Fig. S6.**
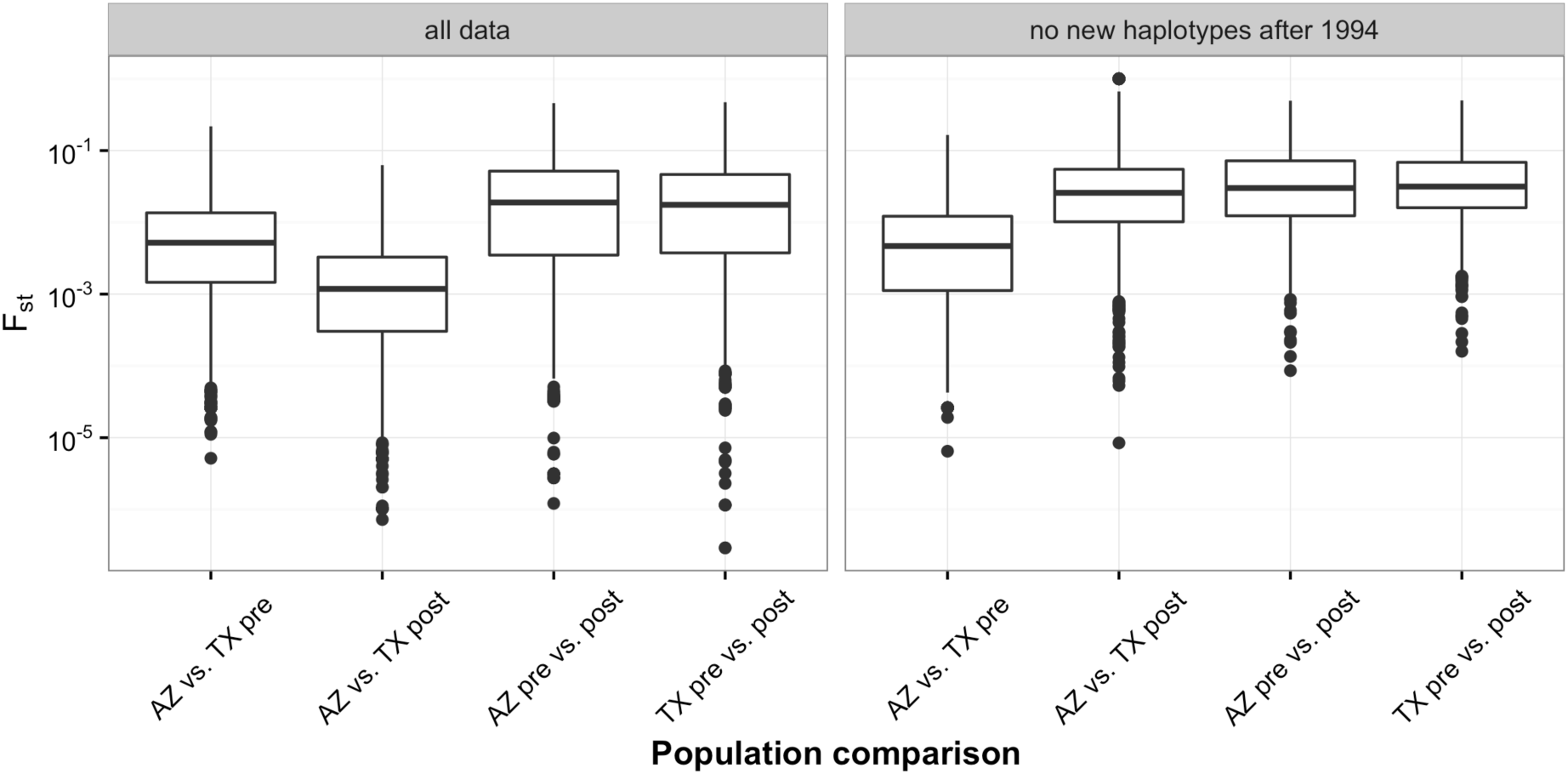
Genetic distances between population pairs for all data (left) and for haplotypes only present before the immigration of Africanized honey bees (right). Consistent with the Africanization process in Arizona and Texas, the post-selection populations had generally lower F_st_ scores (left panel). However, this effect disappeared when immigrant haplotypes were removed (right panel). One data point with a value less that 10^−13^, removed from the AZ vs. TX post comparison in the right-hand plot for clarity.

**Fig. S7.**
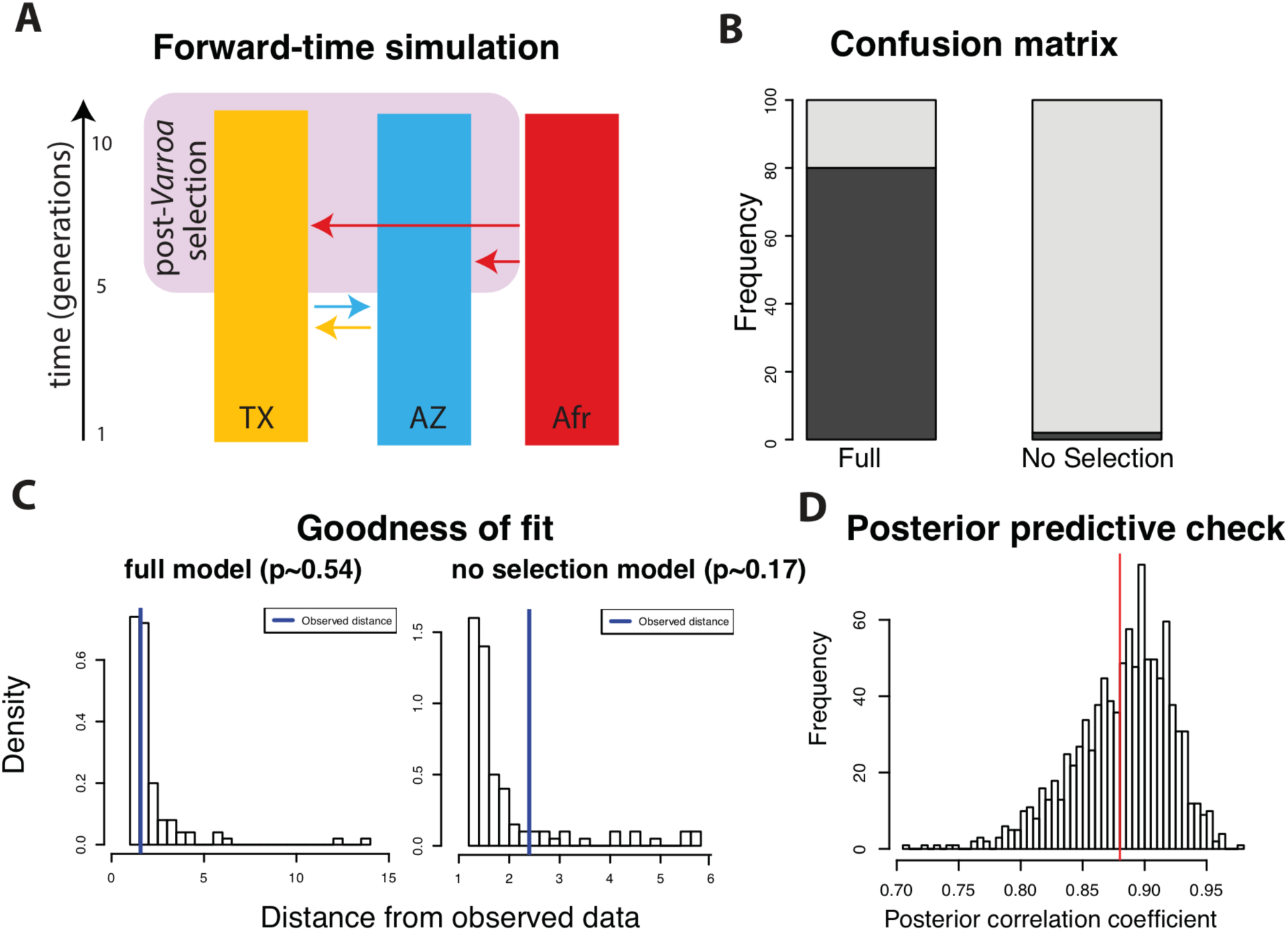
Comparing alternative evolutionary scenarios using Approximate Bayesian Computation. (A) Graphical overview of the model parameters. The “full” and “no selection” scenarios differed in whether the Arizona and Texas populations experienced selection after the arrival of Varroa (pink box). The simulation explicitly modelled observed population reductions (Fig. 1), and estimated migration parameters indicated by arrows, as well as initial population sizes. The rest of the plots show model diagnostics. (B) The confusion matrix indicates that the two evolutionary scenarios could be, in principle, distinguished based on the chosen summary statistic. (C) Goodness of fit could not be rejected for either model, suggesting that they both produce plausible scenarios based on observed data. (D) Posterior predictive check confirms that obtained parameter estimates produce results consistent with observed data. In summary, the two models are technically sound, and have the power to distinguish between alternative evolutionary scenarios.

**Fig. S8.**
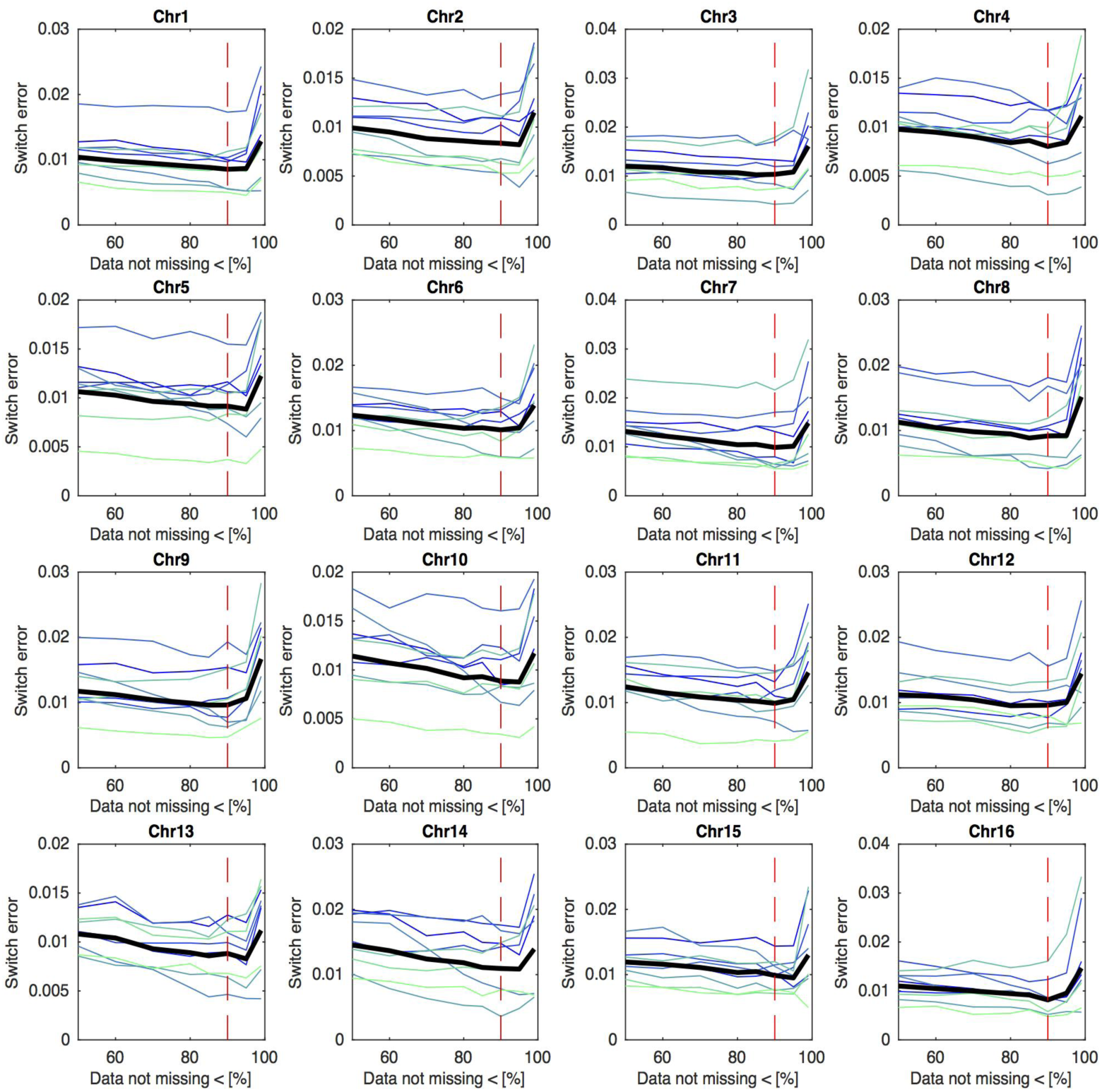
Switch error of the phasing procedure. Switch error change depending on different cut-offs allowing for at most 50, 40, 30, 20, 15, 10, 5, or 1% of missing sequence data in diploid samples for any given position. Switch error was estimated using synthetic diploid drones created by combining together pairs of haploid drone variants. The panels show the switch error values in all 16 honey bee chromosomes. Individual drone results are represented by thin blue lines, while the mean error of all individual synthetic drones are indicated by thick black lines. Vertical red lines indicate the chosen cut-off point of 90% of sequence data detected resulting in the lowest error across the chromosomes.

**Fig. S9.**
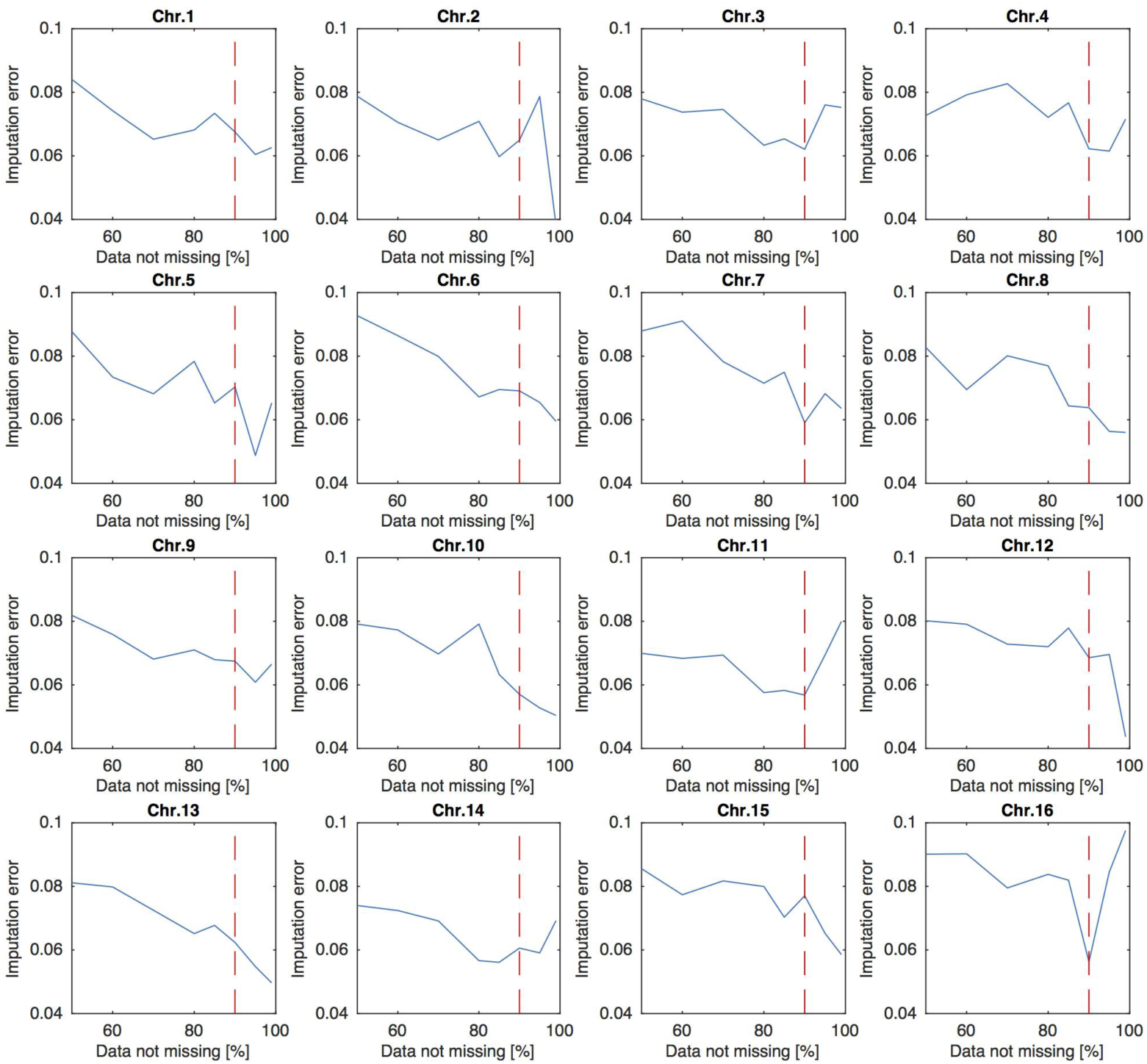
Imputation error. Imputation error change depending on different cutoffs allowing for at most 50, 40, 30, 20, 15, 10, 5, or 1% of missing sequence data in diploid samples for any given position. Imputation error was calculated through comparison of the matching phased and unphased vcf file content of the synthetic diploid drones created by combining together pairs of haploid drone variants. Vertical red lines represent the chosen cut-off point of 90% of sequence data detected resulting in the lowest error across the chromosomes.

**Fig. S10.**
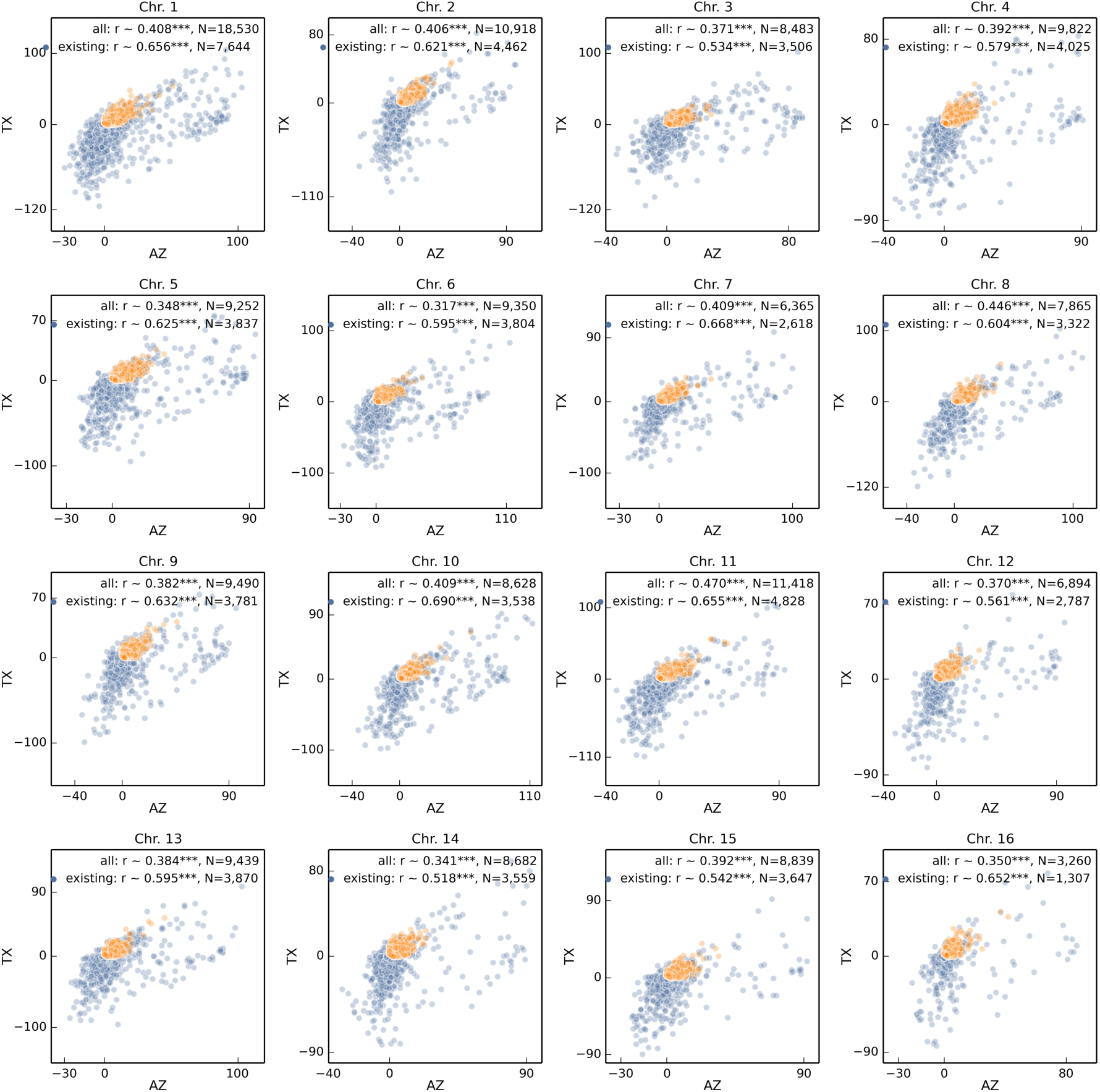
Haplotype count change across chromosomes. Analogous to Fig. 3 correlation of haplotype change between Arizona and Texas populations split among 16 chromosomes.

### Supplemental Tables

**Table S1.**
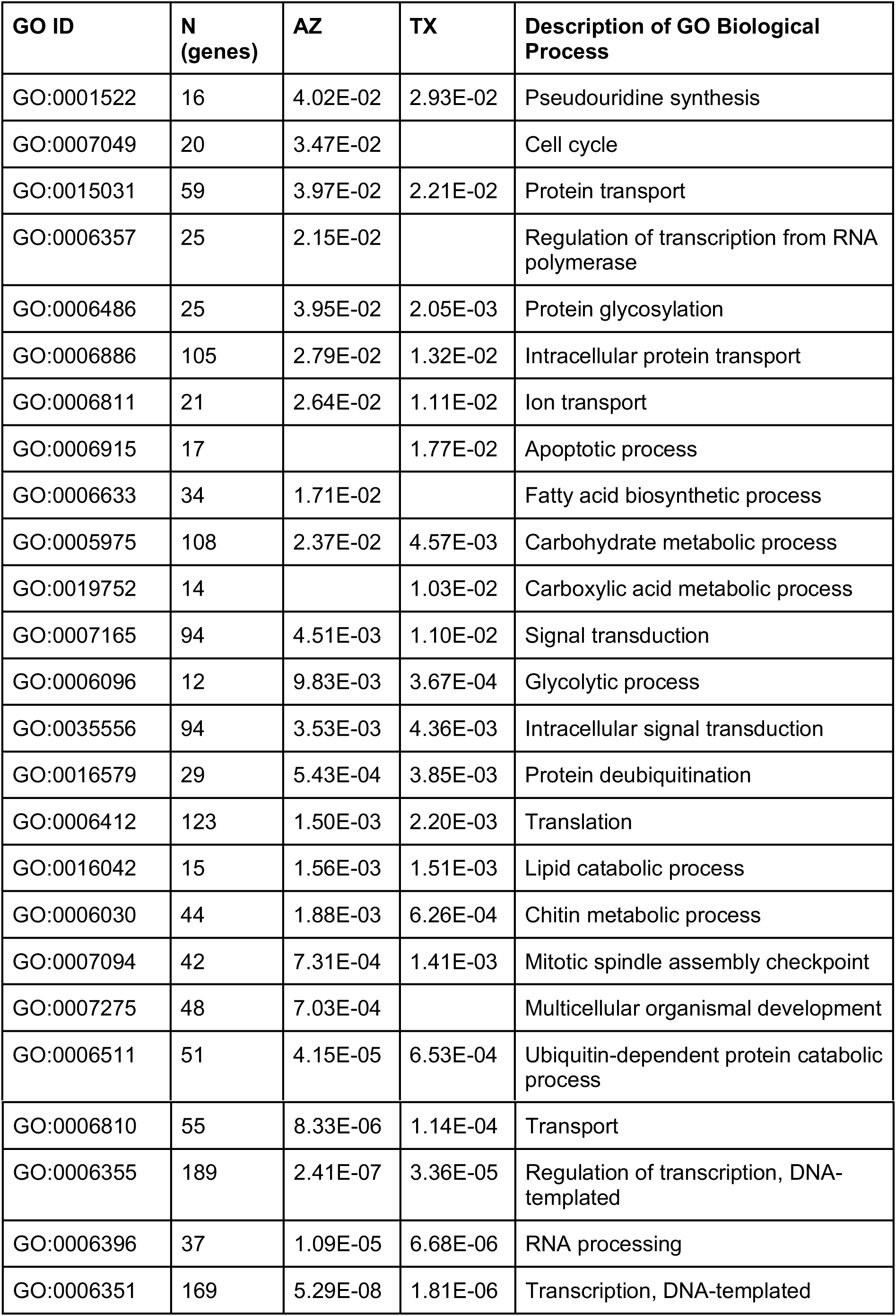
Biological process Gene Ontology (GO) terms significantly associated with changes in haplotype frequency (F_st_) before and after *Varroa* was detected in both populations. Partial regression coefficients quantify the relative contribution of each GO term to the observed change in F_st_ in each population. Coefficients with bootstrap confidence intervals overlapping zero are omitted in the table to reduce visual clutter. Overall GO term partial regression coefficients were highly correlated between Arizona and Texas populations (r = 0.82, d.f. = 199, p < 2 × 10^−16^).

| GO ID | N (genes) | AZ | TX | Description of GO Biological Process |
| --- | --- | --- | --- | --- |
| GO:0001522 | 16 | 4.02E-02 | 2.93E-02 | Pseudouridine synthesis |
| GO:0007049 | 20 | 3.47E-02 |  | Cell cycle |
| GO:0015031 | 59 | 3.97E-02 | 2.21E-02 | Protein transport |
| GO:0006357 | 25 | 2.15E-02 |  | Regulation of transcription from RNA polymerase |
| GO:0006486 | 25 | 3.95E-02 | 2.05E-03 | Protein glycosylation |
| GO:0006886 | 105 | 2.79E-02 | 1.32E-02 | Intracellular protein transport |
| GO:0006811 | 21 | 2.64E-02 | 1.11E-02 | Ion transport |
| GO:0006915 | 17 |  | 1.77E-02 | Apoptotic process |
| GO:0006633 | 34 | 1.71E-02 |  | Fatty acid biosynthetic process |
| GO:0005975 | 108 | 2.37E-02 | 4.57E-03 | Carbohydrate metabolic process |
| GO:0019752 | 14 |  | 1.03E-02 | Carboxylic acid metabolic process |
| GO:0007165 | 94 | 4.51E-03 | 1.10E-02 | Signal transduction |
| GO:0006096 | 12 | 9.83E-03 | 3.67E-04 | Glycolytic process |
| GO:0035556 | 94 | 3.53E-03 | 4.36E-03 | Intracellular signal transduction |
| GO:0016579 | 29 | 5.43E-04 | 3.85E-03 | Protein deubiquitination |
| GO:0006412 | 123 | 1.50E-03 | 2.20E-03 | Translation |
| GO:0016042 | 15 | 1.56E-03 | 1.51E-03 | Lipid catabolic process |
| GO:0006030 | 44 | 1.88E-03 | 6.26E-04 | Chitin metabolic process |
| GO:0007094 | 42 | 7.31E-04 | 1.41E-03 | Mitotic spindle assembly checkpoint |
| GO:0007275 | 48 | 7.03E-04 |  | Multicellular organismal development |
| GO:0006511 | 51 | 4.15E-05 | 6.53E-04 | Ubiquitin-dependent protein catabolic process |
| GO:0006810 | 55 | 8.33E-06 | 1.14E-04 | Transport |
| GO:0006355 | 189 | 2.41E-07 | 3.36E-05 | Regulation of transcription, DNA-templated |
| GO:0006396 | 37 | 1.09E-05 | 6.68E-06 | RNA processing |
| GO:0006351 | 169 | 5.29E-08 | 1.81E-06 | Transcription, DNA-templated |

**Table S2.** KEGG pathways significantly associated with changes in haplotype frequency (F_st_) before and after *Varroa* was detected in both populations. Overall KEGG pathway partial regression coefficients were highly correlated between Arizona and Texas populations (r = 0.84, d.f. = 105, p < 2 × 10^−16^). See also Table S1.

| KEGG ID | N (genes) | AZ | TX | Pathway Description |
| --- | --- | --- | --- | --- |
| ame04214 | 24 |  | 4.8E-02 | Apoptosis |
| ame04745 | 14 |  | 4.7E-02 | Phototransduction |
| ame00190 | 72 | 4.2E-02 |  | Oxidative phosphorylation |
| ame04213 | 21 | 3.6E-02 |  | Longevity regulating pathway |
| ame00020 | 18 | 4.7E-02 | 2.5E-02 | Citrate cycle (TCA cycle) |
| ame04120 | 46 | 4.2E-02 | 2.3E-02 | Ubiquitin mediated proteolysis |
| ame03008 | 44 | 2.8E-02 | 2.4E-02 | Ribosome biogenesis in eukaryotes |
| ame04931 | 15 | 2.7E-02 | 2.2E-02 | Insulin resistance |
| ame04141 | 71 | 1.3E-02 | 1.1E-02 | Protein processing in endoplasmic reticulum |
| ame04146 | 35 | 2.4E-03 | 1.8E-02 | Peroxisome |
| ame04142 | 26 | 8.1E-03 |  | Lysosome |
| ame00480 | 16 | 7.8E-03 |  | Glutathione metabolism |
| ame03013 | 71 |  | 7.2E-03 | RNA transport |
| ame03040 | 66 | 3.8E-03 | 8.1E-03 | Spliceosome |
| ame00564 | 23 | 9.3E-04 | 5.3E-03 | Glycerophospholipid metabolism |
| ame04391 | 19 | 3.5E-03 | 6.6E-04 | Hippo signaling pathway |
| ame04330 | 10 | 5.2E-06 | 4.0E-03 | Notch signaling pathway |
| ame03015 | 35 | 2.5E-03 | 7.5E-05 | mRNA surveillance pathway |
| ame03010 | 98 | 1.3E-04 | 8.0E-06 | Ribosome |
| ame00240 | 33 | 2.6E-07 | 5.8E-05 | Pyrimidine metabolism |
| ame03020 | 19 | 1.3E-08 | 3.9E-06 | RNA polymerase |
| ame00230 | 50 | 1.9E-08 | 3.4E-07 | Purine metabolism |
| ame01100 | 383 | 5.4E-10 | 8.7E-09 | Metabolic pathways |

